# Revisiting the significance of keratin expression in complex epithelia

**DOI:** 10.1101/2022.07.27.501769

**Authors:** Erez Cohen, Craig Johnson, Catherine J. Redmond, Raji R. Nair, Pierre A. Coulombe

## Abstract

A large group of keratin genes (n=54 in the human genome) code for intermediate filament (IF)-forming proteins and show differential regulation in epithelial cells and tissues. Keratin expression can be highly informative about the type of epithelial tissue, differentiation status of constituent cells, and biological context (e.g., normal vs. diseased settings). The foundational principles underlying the use of keratin expression to gain insight about epithelial cells and tissues primarily originated in pioneering studies conducted in the 1980s. The recent emergence of single cell transcriptomics provides an opportunity to revisit these principles and gain new insight into epithelial biology. Re-analysis of single cell RNAseq data collected from human and mouse skin has confirmed long-held views regarding the quantitative importance and pairwise regulation of specific keratin genes in keratinocytes of surface epithelia. Further, such analyses confirm and extend the notion that changes in keratin gene expression occur gradually as progenitor keratinocytes commit to and undergo differentiation, and challenge the prevailing assumption that specific keratin combinations reflect a mitotic vs. a post-mitotic, differentiating state. Our findings provide a blueprint for similar analyses in other tissues, and warrant a more nuanced approach in the use of keratin genes as biomarkers in epithelia.

## Main text

Keratin genes and proteins serve as unparalleled markers towards the identity and status of epithelial cells throughout the body. This reality has been known and exploited for over 40 years (Moll 1982; Fuchs et al. 1985; O’Guin 1990; Fuchs 1995; Moll et al. 2008). Whole genome sequencing efforts in the last two decades expanded our knowledge of the extent of keratin gene diversity and their utility as epithelial markers (Hesse et al. 2004; Schweizer 2006). In humans, 28 type I and 26 type II intermediate filament (IF) genes code for keratin proteins of sizes ranging between 40 and 70 kDa (Schweizer 2006). The dual nature of keratin IF genes (Fuchs 1981) reflects a strict requirement for heteropolymerization of keratin proteins into 10 nm IFs (Coulombe and Fuchs 1990; Hatzfeld and Weber 1990; Steinert 1990). Accordingly, epithelial cells must coordinate the expression of (at least) one type I and (at least) one type II gene to assemble an IF network in their cytoplasm (Sun et al. 1983; Fuchs et al. 1985; Fuchs 1995; Kim and Coulombe 2007). Pioneering studies, largely conducted in the 1980s, established that many type I and II keratin genes are tightly regulated in a pairwise fashion (Sun et al. 1983; Fuchs et al. 1985; Cooper and Sun 1986; Eichner et al. 1986). Further, these pioneering studies revealed the specificity of keratins genes and proteins for the type of epithelial cells, their differentiation status, and whether they are engaged in an adaptative or disease processes (Sun et al. 1983; Fuchs et al. 1985; O’Guin 1990; Fuchs 1995; Moll et al. 2008). Relative to internal “simple” epithelia that occur in the gut, liver, kidney and pancreas (Omary 2017), surface epithelia such as those lining up the skin, oral mucosa and cornea exhibit a greater complexity in keratin expression (Fuchs and Green 1980; Fuchs 1995; Moll et al. 2008). The defining features of the keratin gene family, including their transcriptional regulation and the primary structure of their protein products, are highly conserved in terrestrial mammals (Ehrlich et al. 2019), emphasizing their usefulness and significance as epithelial biomarkers. Accordingly, the signature elements that typify keratin genes and proteins continue to be utilized by researchers and clinicians to interpret epithelial phenotypes when studying disease samples and relevant experimental models (Moll et al. 2008; Rashmi et al. 2009; Karantza 2011; Sharma et al. 2019).

### Revisiting the significance of keratin expression through single cell genomics

The increasing accessibility of single cell RNAseq (scRNAseq) datasets provides a powerful opportunity to revisit the fundamental principles that have been assumed and relied on when using keratin expression to monitor epithelial cell identity or status. In this text, using the epidermis of both human and mouse as examples, we took advantage of available scRNAseq data and paid special attention to three popular notions regarding keratin regulation in surface epithelia; first, that keratins are amongst the most highly expressed genes in keratinocytes; second, that specific pairings of type I and II keratin genes indeed show tight co-regulation in progenitor and differentiating keratinocytes; and third, that specific cell states and localization within epidermis are sharply reflected in keratin expression.

Human trunk skin features a low density of epithelial appendages (hair, glands) along with a relatively thin, low complexity epidermis (Fig. 1A), facilitating an assessment of these principles. We reanalyzed a dataset from human trunk skin reported in Cheng et al. (Cheng et al. 2018) because it features a robust representation of living keratinocytes from across the epidermal sheet (Suppl. Fig. 1A). A total of 24,979 keratinocytes from this study were included in our analyses (cf. Supplemental Material). Given the strategic importance of transgenic mouse models to study epithelial biology in health and disease, we also analyzed a dataset from mouse ear epidermis produced by Lukowski et al. (Lukowski et al. 2018) (Suppl. Fig. 3A). A total of 4,761 keratinocytes from the Lukowski study were included in our analyses (cf. Supplemental Material).

**Figure 1.**
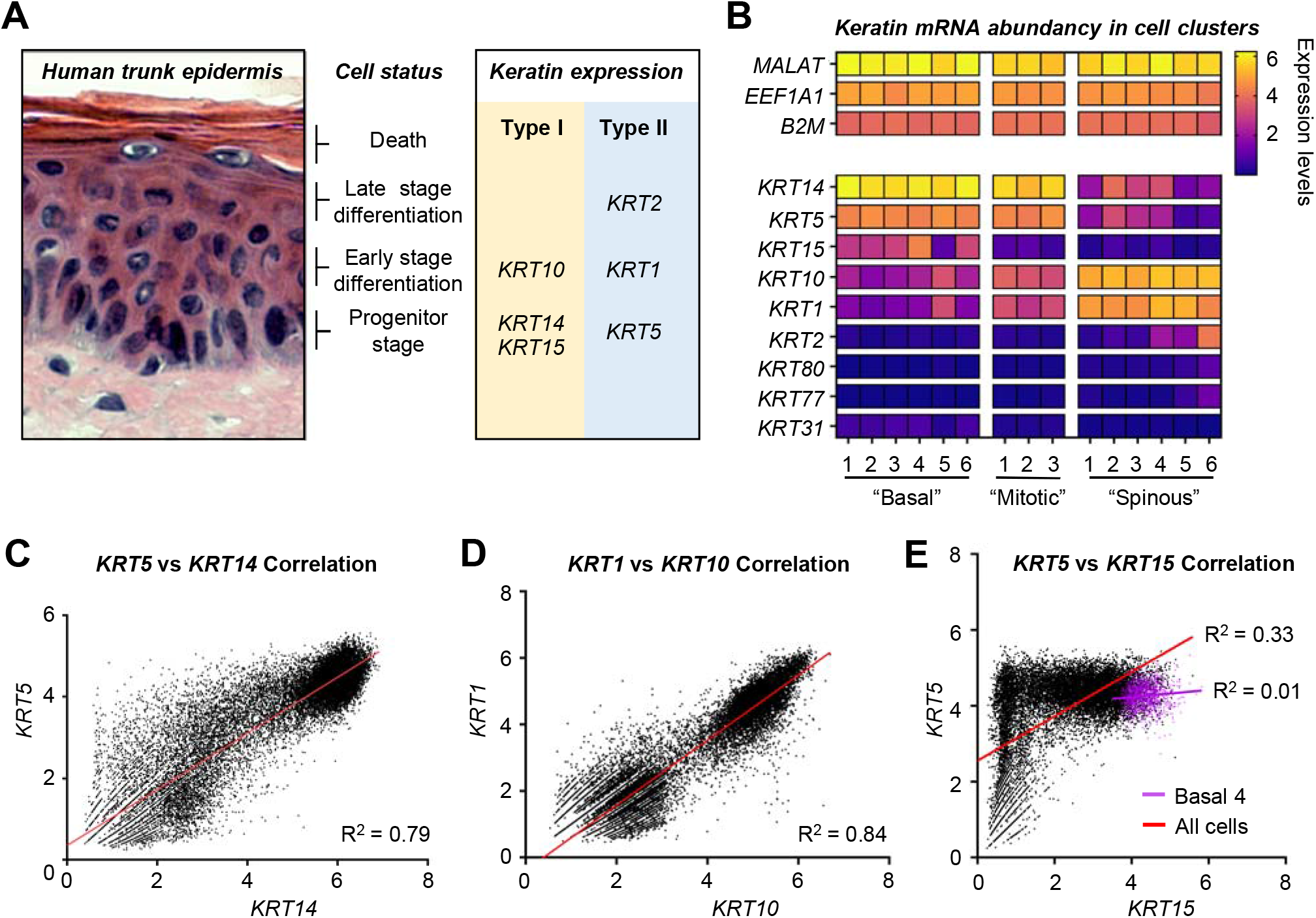
Abundant and pairwise expression of keratin genes in human trunk skin. (A) Cryosection of healthy human trunk skin, stained with hematoxylin-eosin. The status of keratinocytes and their corresponding keratin expression are indicated at right. (B) Heatmap of keratin gene expression in the fifteen Seurat clusters of keratinocytes in healthy human trunk skin, analyzed by sc-RNAseq (Basal1-6; Mitotic1-3, Spinous1-6). Top genes (*MALAT, EEF1A1,B2M*) represent high-expressed genes across all clusters. (C-E) Correlations between specific type I and type II 2 keratin genes across the entire array of epidermal keratinocytes (n= 24,979 cell). Red lines represent linear correlations, and R^2^ values are reported. (C) *KRT5* vs *KRT14*. (D) *KRT10* vs *KRT1*. (E) *KRT5* vs *KRT15*. In E, the magenta line conveys the linear correlation for the Basal4 cluster, specifically.

### Abundance of keratin gene expression in normal human interfollicular epidermis

Keratinocytes of conventional interfollicular epidermis primarily express six keratin genes. These are the type II IF *KRT5* and the type I IFs *KRT14* and *KRT15*, characteristic of progenitor cells located in the basal layer (Fuchs and Green 1980; Nelson and Sun 1983; Lloyd et al. 1995), the type II *KRT1* and type I *KRT10*, characteristic of early stage differentiating cells (Fuchs and Green 1980; Woodcock-Mitchell et al. 1982), and the type II IF *KRT2*, characteristic of late stage differentiating cells (Collin 1992; Collin et al. 1992) (Fig. 1A). The number of reads for a given gene, when normalized to library size, directly relates to abundance of mRNA transcripts in that given cell in a scRNAseq data set. *MALAT* (Wu et al. 2015), *EEF1A1* (Abbas et al. 2015) and *B2M* (Li et al. 2016) are examples of genes for which the number of reads is consistently at the top across most cell clusters in Cheng et al. (Cheng et al. 2018) (Fig. 1B), thus providing a reference for highly expressed genes in human epidermis.

In relatively undifferentiated keratinocytes (e.g., “Basal1-6” clusters; Suppl. Fig. 1A), *KRT5* and especially *KRT14* are among the highest abundance mRNAs in human trunk skin epidermis (Fig. 1B and Suppl. Table 1). Relative to these *KRT5* and *KRT14*, the *KRT15* mRNA is more restricted in its expression domain and expression levels (Fig. 1B). *KRT15* is one of the ten most expressed genes in the “Basal4” cluster (Suppl. Table 1), which is populated by cells exhibiting a strong basal lamina gene signature (*POSTN, DST, COL17A1*), high levels of *KRT14* mRNA (Suppl. Tables 1 and 2), and are predominantly in the G1 phase of the cell cycle (Suppl. Fig. 1, D and E). Across all relevant clusters, the average number of reads per cell for the type I *KRT14*, either alone or combined with *KRT15*, are markedly higher than those of *KRT5*, the sole type II keratin gene expressed in undifferentiated keratinocytes of interfollicular epidermis (Suppl. Table 2, Suppl. Fig. 2A). These observations also apply to mitotic keratinocytes (“Mitotic1-3” clusters; see Fig. 1B, Suppl. Tables 1 and 2). Since the size of mature mRNAs is similar for type I (<2.0 kb) vs. type II keratins (<2.5 kb) (Fuchs 1981; Roop et al. 1983), size-related biases likely do not contribute to differences in transcript abundance.

In keratinocytes undergoing differentiation (e.g., “Spinous1-6” clusters; Suppl. Fig. 1A), *KRT1* and *KRT10* represent highly abundant transcripts (Fig. 1B and Suppl. Table 1). In this instance, the number of reads for *KRT1* and *KRT10* transcripts is closer to a 1:1 ratio (Suppl. Fig. 2A and Suppl. Table 2). The *KRT2* mRNA is considerably more restricted in both its expression domain and levels relative to each of *KRT1* and *KRT10* (Fig. 1B and Suppl. Table 1). *KRT2* is one of the ten most expressed genes in only one cluster, “Spinous6”, which is populated by cells that express late differentiation markers (Suppl. Table 1) and in G1 phase of the cell cycle (Suppl. Fig. 1, D and E).

These scRNAseq analyses confirm the notion that the six keratin genes known to be expressed at significant levels in keratinocytes of the epidermis, namely, the type II IF *KRT1, KRT2*, and *KRT5* and the type I IF *KRT10, KRT14*, and *KRT15*, are among the most highly expressed genes in cells making up this epithelium.

### Pairwise regulation of keratin genes in human trunk epidermis

The Cheng et al. dataset (Cheng et al. 2018) also provides a straightforward opportunity to re-examine pairwise regulation of type I and II keratin genes in healthy human epidermis. Along with the *KRT8-KRT18* pair emblematic of simple epithelia (Omary 2017), *KRT5-KRT14* and *KRT1-KRT10* are understood to be among the tightest keratin pairings in all of epithelial biology. Expression of keratin pairs at a single cell level was plotted across all epidermal keratinocytes, totaling 24,979 cells, and linear correlations in keratin expression across cells and cell clusters in human trunk skin were assessed. Very high Pearson correlation coefficients occur between *KRT5* and *KRT14* mRNA reads (r^2^= 0.79; p<0.0001; Fig. 1C) and *KRT1* and *KRT10* mRNA reads (r^2^= 0.84; p<0.0001; Fig. 1D). We proceeded to test if similar conclusions apply when considering relevant individual clusters (Suppl. Table 3). A reduction in statistical robustness is expected for such analyses due to smaller cell numbers in single clusters along with assessing correlations in restricted ranges such as defined clusters (Bland and Altman 2011). Regardless, *KRT1* and *KRT10* are highly correlated in the “Spinous4” cluster (n=1007 cells; r^2^= 0.82; p<0.0001), which features the highest number of reads for both genes. Similarly, *KRT5* and *KRT14* show a lesser though significant correlation in, e.g., the “Basal1” cluster (n=5037 cells; r^2^= 0.35; <0.0001), which shows the highest levels of *KRT14* mRNA reads (Suppl. Table 3). The degree to which *KRT5* and *KRT14*, and *KRT1* and *KRT10*, are each coordinately regulated is truly remarkable when factoring in the notion that type I and type II keratin genes are clustered on separate chromosomes in the human genome and that of other mammals (Powell et al. 1986; Lessin et al. 1988; Romano 1988). As of yet, there is no mechanistic basis for the co-regulation of type I and II keratin genes in either general or specific terms.

Keratin gene-specific correlations are far less compelling for the more restricted *KRT15* (type I; progenitor keratinocytes) and *KRT2* (type II; differentiating keratinocytes). Across the entire array of keratinocytes in human trunk skin, *KRT15* and *KRT5* (type II) exhibit a markedly lower Pearson correlation coefficient (r^2^ = 0.33) than *KRT5-KRT14* (Fig. 1E). Surprisingly, *KRT15* exhibits a higher Pearson correlation coefficient with type I *KRT14* (r^2^ = 0.48) (Suppl. Table 3). Importantly, high expression of two keratins does not necessarily imply linear correlation at a single-cell level. For instance, linear regression analysis indicates no correlation between the two genes (r^2^ = 0.01) in the cell cluster featuring the highest expression of *KRT15* and *KRT5* (“Basal4”; Fig. 1E and Suppl. Table 3). For *KRT2*, the correlation observed with type I *KRT10* also appears relatively weak, given r^2^ = 0.14 for trunk skin cells as a whole. In this case, the two genes show a stronger correlation in the cluster that expresses *KRT2* at the highest level (“Spinous6”, r^2^ = 0.56; Suppl. Fig. 2B and Suppl. Table 3). From these findings, we infer that there are keratin genes, e.g., *KRT15*, for which pairwise regulation does not represent a compelling attribute at the mRNA transcript level, at least in human trunk skin. It will prove interesting to test whether this notion applies to other epithelia in which *KRT15* transcript levels are higher relative to KRT14 and/or other type I keratins.

### Emerging concept in keratinocyte biology: hybrid expression of progenitor- and differentiation-related keratins

The transition that occurs between progenitor and differentiating keratinocytes in interfollicular epidermis is considered by many to be sharp and tighly coupled to the transition between *KRT5-KRT14* and *KRT1-KRT10* expression. Onset of *KRT1* and *KRT10* expression in healthy interfollicular epidermis is widely believed to occur when progenitor keratinocytes in the basal compartment become post-mitotic and enter the suprabasal compartment. However, this notion does not accurately reflect foundational studies on keratin expression published in the 1980s (Fuchs and Green 1980; Schweizer et al. 1984; Regnier et al. 1986; Roop 1987). Such views on the respective significance of *KRT5*-*KRT14* and *KRT1*-*KRT10* expression originate, in part, from the spatial staining patterns afforded by several (but not all) antibodies to these keratin proteins in immunostained tissue sections under standard imaging conditions (e.g., “suprabasal” K14 is often equated with hyperproliferation in the literature). Indeed, combining experimental evidence with technical advances in sequencing, a growing body of literature reports on keratinocytes with hybrid features including the expression of progenitor- and differentiation-related keratins. Initial observations of these populations were established in 2020 in both newborn mice (Lin et al. 2020) and neonatal human foreskin (Wang et al. 2020) where sc-RNAseq data identified hybrid interfollicular epidermal transition cells. While both studies utilized experimental methods to validate these findings *in-vivo* (RNA-FISH (Lin et al. 2020) and immunohistochemistry (Wang et al. 2020)), they differed in the observed proportion of transitional cells in the dataset (with 20% observed in newborn mouse skin vs. 4% in human neonatal foreskin). While these discrepancies could reflect differences on a tissue or organismal level (which our analysis supports), we note that the exact proportion of these hybrid cells among all keratinocytes could be biased due to methodological differences in cell dissociation methodology, impacting keratinocyte populations sequenced and the heterogeneity reflected (Kim et al. 2020). Additional studies have since utilized immunohistochemistry to observe these transitional cells, and identify them as basal cells committed for differentiation (*Krt14*+*Krt10*+), yet capable of cycling (Aragona et al. 2020). Utilizing a pulse-chase BrdU approach Aragona et al. observed that 36% of cycling cells in mouse dorsal skin were positive for both K14 and K10, suggesting that such transient cells might still actively cycle under homeostatic conditions (Aragona et al. 2020). Most recently, this paradigm was further explored under live-imaging *in-vivo* utilizing lineage tracing and *Krt10* expression reporter (Cockburn 2021). Further supporting the existence of a cycling, transient population, the authors were able to follow basasl cells commitment to differentiation under live imaging, noting the induction of *Krt10*, and observing proliferation of these hybrid, differentiation-commited cells (Cockburn 2021). With these observations in mind, we wanted to identify whether an approach focused on keratin regulation supports these emerging findings, as well as identify whether such population might be conserved in the human trunk epidermis. We set out to examine these issues anew through a multi-pronged approach that included i) relating *KRT14* and *KRT10* expression in the 24,979 single epidermal keratinocytes from the Cheng et al. (Cheng et al. 2018) human data set and ii) analyze this relationship from the perspective of genes that are reliably associated with the G2/M phase of the cell cycle and localization to the basal layer of epidermis.

Relating *KRT10* to *KRT14* in a scatter plot (Fig. 2A) reveals that large numbers of epidermal keratinocytes partition to a *KRT14*^high^/*KRT10*^low^ subgroup (53% of the pool), consistent with a progenitor character, or to a *KRT14*^low^/*KRT10*^high^ subgroup (37% of the pool), consistent with a differentiating character. The observation that *KRT14*^high^/*KRT10*^low^ represent a larger group of cells in the sc-seq data is not surprising and could be due to methodological ease of obtaining single cells from the basal rather than the suprabasal differentiating compartment of epidermis (Kim et al. 2020; Burja et al. 2022). However, we note that the dataset contains a large fraction of cells expressing late-stage differentiation markers (such as *KRT2*; Fig. 1B), and that 37% of the keratinocyte pool represents 9,230 cells, indicative of the robustness of the dataset in capturing subpopulations of keratinocytes at varying stages of differentiation. With this caveat alleviated to a significant degree we find that, surprisingly, a sizable fraction of keratinocytes express appreciable levels of both *KRT14* and *KRT10* transcripts (9.5% of the pool). That such *KRT14*^high^/*KRT10*^high^ keratinocytes exhibit a hybrid character is further substantiated by their content in mRNAs associated with both a progenitor state (e.g., *POSTN, DST, COL17A*) and a differentiated state (e.g., *DMKN, PSP, KRTDAP, SBSN*). In contrast, no cells with hybrid characteristics were identified when *KRT15* and *KRT2* levels were related in such a scatter plot (Suppl. Fig. 2C).

**Figure 2.**
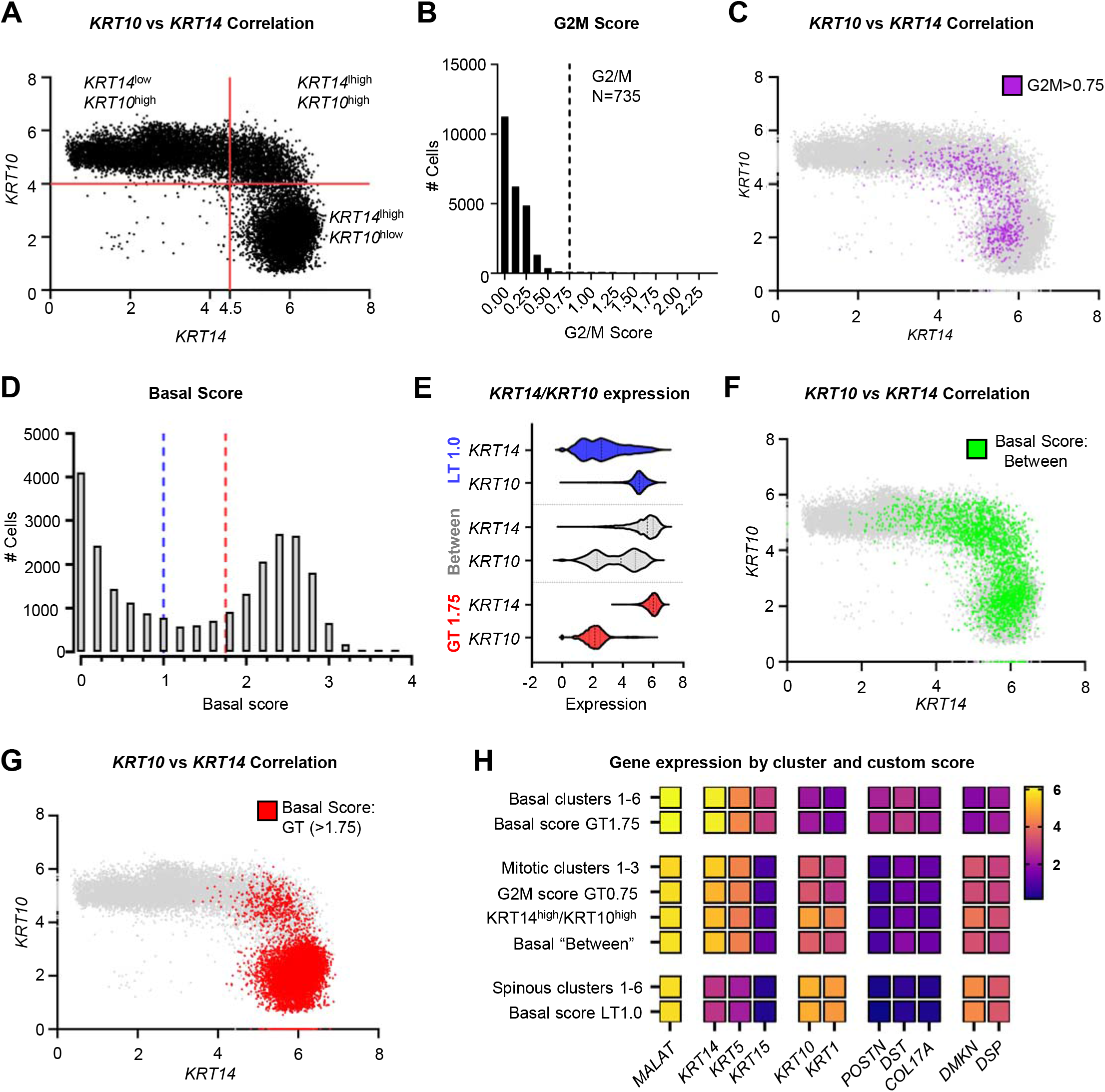
scRNA-seq data identifies a *KRT10*^*high*^/*KRT14*^*high*^ hybrid keratinocyte population. (A) Correlation of *KRT10* and *KRT14* in human trunk epidermis (n=24,979 cells). Quadrants were defined by abundancy of cells expressing high *KRT14* (>4.5) or high KRT10 (>4.0). (B) Histogram reporting on the composite G2/M score across the same cell population. Dashed line represent threshold for G2/M cells (>0.75, ∼3% of cells). (C) Mapping of G2/M cells with a high (>0.75) G2/M score (pink) on the *KRT10*-*KRT14* scatter plot show in frame A. (D) Histogram reporting on the composite “Basal Score” across the population of epidermal keratinocytes. Blue and red dashed lines partition the cells into low (lower than 1.0 or LT1.0), high (greater than 1.75 or) and “Between” Basal Score groups. (E) Violin plots of *KRT14* and *KRT10* expression in cells with LT1.0, “Between”, and GT1.75 Basal Scores. (F-G). Mapping of cells with a (F) “between” and a (G) GT1.75 basal score on the *KRT10*-*KRT14* scatter plot shown in frame A.(H) Heatmap representation of expression levels for genes associated with basal and spinous differentiation state in Seurat clusters and custom scores.

Among the myriad of questions that arise about the KRT14^high^/KRT10^high^ keratinocytes with hybrid characteristics, two that stand out relate to their mitotic competency and location within the trunk epidermis. To examine the mitotic state of keratinocytes exhibiting hybrid features, we devised a combined G2/M score based on average expression of five genes that are highly specific to this phase of the cell cycle: *MKI67* (Booth et al. 2014), *TOP2A* (Earnshaw et al. 1985), *CDC20* (Kramer et al. 2000), *BUB3* (Kalitsis et al. 2000), and *CCNB2* (Nigg 2001). The rationale for using these five genes in combination is to stabilize the variation one would expect from a single gene’s data points. Sorting all epidermal keratinocytes from human trunk skin according to this combined G2/M score yielded a left-skewed, strongly asymmetric distribution, as expected (Fig. 2B). To identify keratinocytes that are indisputably mitotic, we applied a high cut-off filter, set at 0.75, yielding 735 out of 24,979 keratinocytes, or 2.94% of the total cell population (Fig. 2B). Cells with such a high G2/M score are almost entirely comprised within the “Mitotic2” cluster in the original UMAP (Suppl. Table 4). Moreover, *KRT14* and *KRT5* are among the highest expressed genes in this subgroup (Fig. 2H). Remarkably, however, so are *KRT10* and *KRT1*, along with other genes previously associated with keratinocyte differentiation including *DMKN* (Bazzi et al. 2007) and *DSP* (Lechler and Fuchs 2007) (Fig. 2H). Mapping keratinocytes showing a high composite G2/M score (GT>0.75) onto the *KRT10*/*KRT14* scatter plot reveals that cells expressing mitotic genes occur in both the *KRT14*^high^/*KRT10*^low^ and *KRT14*^high^/*KRT10*^high^ quadrants (Fig. 2C). These findings suggest the existence of a keratinocytes population that have initiated differentiation, as reflected through expression of *KRT10, KRT1*, and several additional markers (*DMKN, PSP, KRTDAP,SBSN*), yet can maintain a mitotic signature in human trunk skin epidermis.

We next sought to assess the location of keratinocytes with hybrid features in human trunk skin epidermis. A combined basal layer score incorporating the expression levels for three genes encoding proteins specifically involved in anchoring keratinocytes to the basal lamina, *DST* (Sawamura et al. 1991), *POSTN* (Nishiyama et al. 2011), and *COL17A1* (McGrath et al. 1995), was devised. Individually, *DST, POSTN* and *COL17A1* show similar distributions across cells in this data set (Suppl. Fig. 2D). Sorting all keratinocytes (n= 24,979) using the combined basal score yields a classic bimodal distribution, reflecting in part the expected downregulation of basal lamina associated genes within the suprabasal compartment (Fig. 2D). Keratinocytes were next partitioned into three subsets based on the basal score (Fig. 2D). When considering either “top marker genes” or “highest-expressed genes”, keratinocytes exhibiting a high basal score (“GT>1.75”; n= 12,123 cells) are characterized by a strong progenitor signature (e.g., Fig. 2, E and H) consistent with a location within the basal layer of epidermis. On the other hand, keratinocytes with a low basal score (“LT<1.0”; n= 10,402 cells) show a strong differentiation signature when using the same criteria (Fig. 2, E and H), consistent with a location away from the basal compartment. With such expectations met, it proves interesting to analyze the subset of keratinocytes showing an intermediate basal score (“between”; n= 2,454 cells) in this analysis (Fig. 2D). Cells with a intermediate basal score express both *KRT14* and *KRT10* at appreciable levels (Fig. 2E) and populate a broad fraction of keratinocytes as sorted in the *KRT10*/*KRT14* scatter plot analysis (Fig. 2F). Remarkably, a sizable fraction (34%) of keratinocytes with an intermediate (“between”) basal score map to the *KRT14*^high^/ *KRT10*^high^ subgroup in the *KRT10*/*KRT14* scatter plot (Fig. 2F). This enrichment (34% observed vs 18% expected, χ2 = 2,216, p-value < .0001) is striking since *KRT14*^high^/ *KRT10*^high^ hybrid cells populate considerably smaller fractions of both high and low basal scoring groups (e.g., 3.6% of cells within the GT>1.75 subset vs. 10% expected, and 10.2% of cells in LT<1.0 vs. 10% expected). Conversely, 19% of keratinocytes exhibiting a high (GT>1.75) basal score map to the *KRT14*^high^/ *KRT10*^high^ subgroup in the *KRT10*/*KRT14* scatter plot (Fig. 2G).

Taken together, the evidence stemming from analyses of scRNAseq collected from human trunk skin suggests the existence of a sizable subpopulation of keratinocytes that: i) reside in the basal layer; ii) express intermediate levels of *KRT5*-*KRT14, KRT1*-*KRT10*, and additional genes reflecting a differentiating status (Fig. 2H); and iii) can undergo mitosis as reflected by relatively high levels of G2/M gene transcripts. From this we conclude that changes in keratin gene expression occur quite gradually as keratinocytes of the epidermis transit from a progenitor to a differentiating state (see (Stoler 1988), and also that expression of the differentiation-related *KRT1*/*KRT10* can occur in keratinocytes undergoing mitosis in the epidermis. The outcome of this keratin expression-focused compuatational analysis thus confirm and extend the efforts of others combining computational and experimental approaches (Aragona et al. 2020; Lin et al. 2020; Wang et al. 2020; Cockburn 2021).

While our computational analysis focuses on the transcriptional landscape of keratinocytes, it remains important to bridge the gap and reexamine the relation between mRNA and protein levels when studying this emerging concept in keratinocyte biology. Work by our lab and others have attempted to address these questions in the past, measuring cellular protein levels of K5, K14 and K15 in sorted basal keratinocytes (Sun and Green 1978; Feng 2013) (Feng et al, JID 2013, Sun & Green 1978). K5 and K14, in particular, are highly expressed at both the mRNA and protein levels in basal keratinocytes. Further correlating these studies, as well as measurements of K10 protein on a single-cell level would likely prove highly informative in understanding the nature of keratinocyte commitment to differentiation. This is especially important considering evidence for a role of post-translational modifications in keratinocyte differentiation (Guo et al. 2020) and will further establish our defintions of low/high expression of genes/protein and their transitions across keratinocytes differentiation and cycling.

### Key attributes of keratin gene regulation are conserved in mouse ear skin epidermis

Transgenic mouse models have made an enormous contribution to our understanding of gene function and disease mechanisms in human skin epithelia, prompting us to next assess whether the principles emanating from our analysis of human trunk skin scRNAseq data also applies to mouse skin. To do so, we re-analyzed the scRNAseq data set from Lukowski et al. (2018) reporting on intact mouse ear skin from mouse ear skin (Supplemental Material and Suppl. Fig. 3A), which arguably shares many attributes with trunk skin in the human. Focusing on the six cell clusters totaling 4,761 cells that comprise epidermal keratinocytes, we confirmed that *Krt5* and especially *Krt14* reads occur at high levels relative to known abundant transcripts (e.g., *Eef1a1*) in clusters enriched in progenitor keratinocytes (“Basal1,2”; Fig. 3A). Likewise, *Krt1* and especially *Krt10* reads occur at high levels relative to abundant genes in clusters enriched in differentiating keratinocytes (“Spinous1-3”; Fig. 3A). Moreover, *Krt5* and *Krt14* genes are highly correlated across the entire array of 4,761 keratinocytes in mouse ear (r^2^ = 0.74; Fig. 3B) while, again, the degree to which *Krt15* is correlated to *Krt5* is lower (r^2^ = 0.53; data not shown). In this setting, *Krt10* transcripts are far better correlated to those of *Krt2* (r^2^ = 0.35; Fig. 3C) compared to *Krt1* (r^2^ = 0.13; data not shown), a novel and interesting finding that merits follow-up studies. Relating *Krt14* to *Krt10* expression through a scatter plot analysis of all keratinocytes of mouse ear epidermis gives rise to a distribution which, again, includes *KRT14*^high^/*KRT10*^high^ hybrid keratinocytes in addition to the expected *KRT14*^high^/*KRT10*^low^ and *KRT14*^low^/*KRT10*^high^ keratinocyte subsets (Fig. 3D). We also analyzed this cell population through the G2/M composite score, exactly as done for human skin (see Fig. 2B). Filtering the resulting distribution using a high G2/M score (>0.6) yielded 266, or 5.5%, keratinocytes (Fig. 3E) that largely binned to the mitotic cluster group (81%; Suppl. Table 4). In contrast to the human trunk dataset, relatively few keratinocytes with a high G2/M score map to the hybrid keratinocytes in mouse ear skin, given that only 1.4% of *KRT14*^high^/*KRT10*^high^ cells exceed the composite G2/M score threshold (Fig. 3F; by comparison, 8.3% of the high score G2/M cells map to this hybrid group in human trunk skin). Instead, 91% of all high G2/M scoring cells reside in the *KRT14*^high^/*KRT10*^low^ group (Fig. 3F). Whether these differences are due to the lower statistical power and/or missing keratinocyte supopulations in the mouse skin study (Lukowski *et al* 2018), or are related to the reality of distinct body location (human trunk vs. mouse ear), or reflect deeper and more significant biological and/or evolutionary differences between human and mouse skin, remains to be explored.

**Figure 3.**
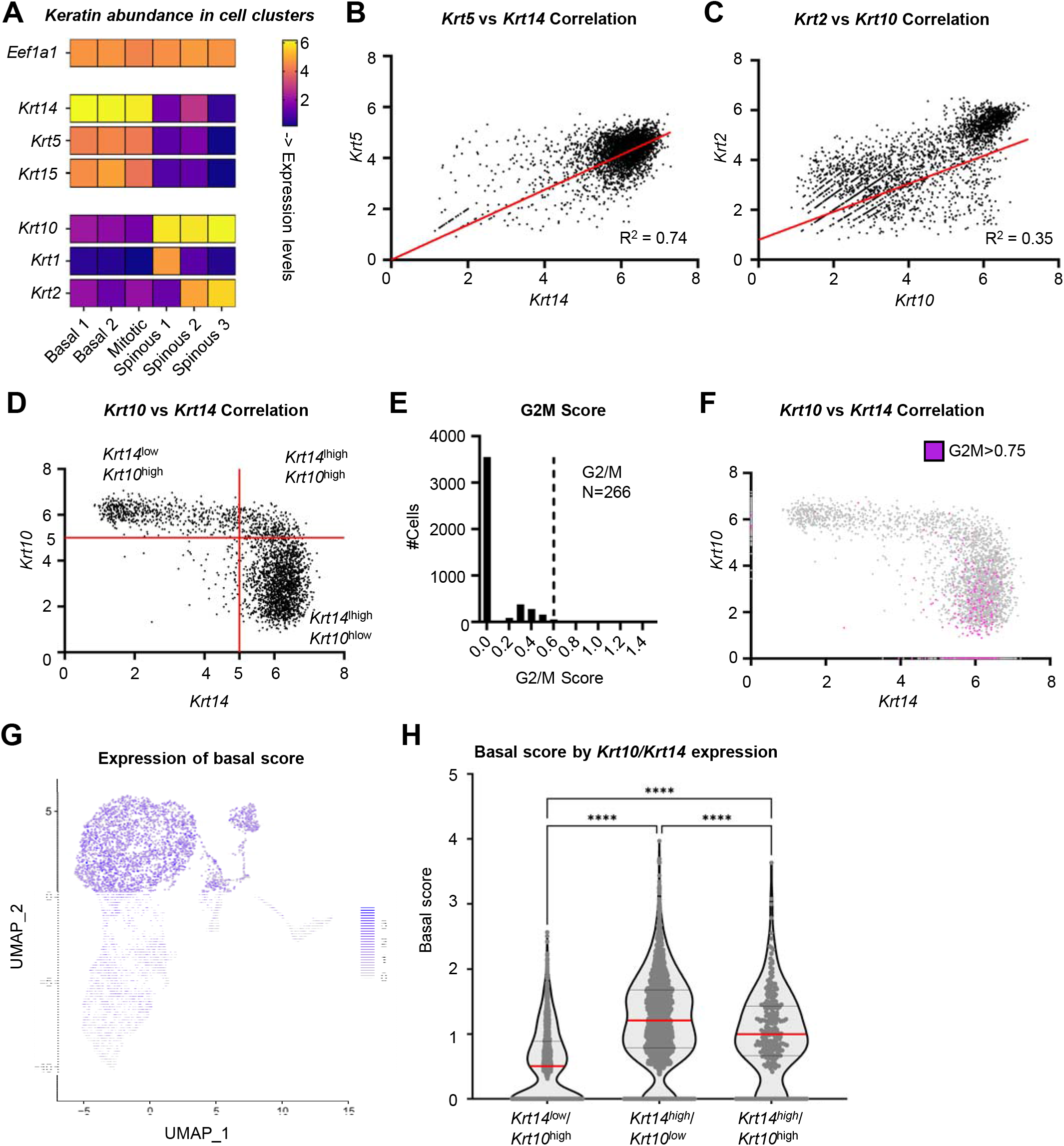
Keratin expression in mouse ear skin. (A) Heatmap of keratin gene expression in the six Seurat clusters of keratinocytes (n=4,761 cells) in healthy mouse ear skin, analyzed by sc-RNAseq (Basal1-2; Mitotic, Spinous1-3). *Eef1a1* provides a reference for a highly expressed gene across all clusters. (B-C) Correlations specific type I and type II 2 keratin genes across the entire array of epidermal keratinocytes in mouse ear skin (n= 4,979 cells). Red lines represent linear correlations, and R^2^ values are reported. (B) *Krt5* vs *Krt14*. (C) *Krt2* vs *Krt10*. (D) Correlation of *Krt10* and *Krt14* in mouse ear epidermis. Quadrants were defined by abundancy of cells expressing high *Krt14* (>5.0) or high Krt10 (>5.0). (E) Histogram reporting on the composite G2/M score across the same cell population. Dashed line represent threshold for G2/M cells (>0.6, ∼5.5% of cells). (F) Mapping of G2/M cells with a high (>0.6) G2/M score (pink) on the *Krt10*-*Krt14* scatter plot shown in frame D. (G) Mapping of the Basal Score for individual cells on the original UMAP from the mouse ear skin data set (see Suppl. Fig. 3). (H) Violin map representation of Basal Score in cells that express low and/or high combinations of *Krt14* and *Krt10* (see frame D). **** represent adj.p <0.0001 as tested by one way ANOVA with Tukey’s multiple comparisons test.

Utilizing basal lamina-anchoring genes (*Postn, Dst, Col17A1*; see Suppl. Fig. 4A) as we did when analyzing the human keratinocyte data set, we compiled an average basal score for each cell and examined the distribution of scores for epidermal keratinocytes across mouse ear skin. The resulting distribution (Suppl. Fig. 4B) did not follow a classic bimodal shape as it did for human trunk skin epidermis, again possibly the result of underrepresentation of differentiating keratinocytes in the data set. Still, plotting the basal score on the original UMAP revealed that keratinocytes with a high basal score (GT>1.8) are contained within the “Basal 1-2” and “Mitotic” cell clusters (93% of cells; Fig. 3, G and H). We then analyzed the distribution of basal score in the setting of a scatter analysis relating *Krt14* and *Krt10* expression. *Krt14*^high^/*Krt10*^low^ cells are defined by an average basal score (1.23 ± 0.01) that is significantly higher than that of *Krt14*^low^/*Krt10*^high^ cells (0.53 ± 0.01, P<0.0001). Importantly, and consistent with our analysis of human trunk skin data, *Krt14*^high^/*Krt10*^high^ hybrid keratinocytes in mouse skin epidermis show an intermediate score (1.0 ± 0.03; Fig. 3H), suggesting that these cells maintain a similar location, and progenitor-like gene expression. Though more analyses are needed, such observations confirm that the key principles governing keratin gene expression in the interfolliclar epidermis of human trunk skin (Figs. 1-2) are conserved in mouse skin (Fig. 3).

### Conclusions and future outlook

The power of single cell RNAseq analyses has exposed a significant molecular heterogeneity within the classically defined ‘progenitor’ and ‘differentiating’ compartments in human and mouse interfollicular epidermis (Joost et al. 2016; Cheng et al. 2018; Lukowski et al. 2018; Aragona et al. 2020; Dekoninck et al. 2020; Haensel et al. 2020; Lin et al. 2020; Wang et al. 2020; Cockburn 2021). Whether this heterogeneity reflects a continuum of cellular states in the setting of a single developmental/differentiation pathway and/or distinct classes of progenitor stem cells and/or co-existence of distinct differentiation pathways within epidermis is a fundamentally important issue that must await further experimentation and consideration (see (Lin et al. 2020; Wang et al. 2020; Cockburn 2021)). Here, we considered specific aspects of keratin expression in interfolliclar epidermis from a perspective of single cell RNAseq analyses, and offer the following three conclusions. First, keratin-encoding genes are among the most abundantly expressed genes in epidermal keratinocytes. Second, the type I *KRT14* and type II *KRT5* and the type I *KRT10* and type II *KRT1* are each tightly co-regulated at the mRNA transcript level in epidermal keratinocytes, independently of the cellular state, thereby justifying a designation of pairwise regulation for them. By contrast, two additional keratin genes expressed at significant levels in the epidermis, i.e., the type II *KRT2* in late stage differentiating keratinocytes and the type I *KRT15* expressed in undifferentiated keratinocytes in the basal layer, do not show co-regulation with a dedicated partner keratin gene of the complementary type, at least at the transcript level. Third, our analyses support recent observations of changes in keratin gene expression occuring while keratinocytes transit from a progenitor to a differentiating state in the interfollicular epidermis, and also that expression of the differentiation-related *KRT1*/*KRT10* occurs in keratinocytes that maintain a mitotic gene expression signature. Though onset of *KRT1*-*KRT10* expression clearly conveys an engagement of keratinocytes towards differentiation, it does not necessarily reflect a post-mitotic state, consistent with early studies in the field (Schweizer et al. 1984; Regnier et al. 1986) and more recent studies also based on high throughput single cell analyses (Lin et al. 2020; Cockburn 2021). We predict that similar conclusions will be attained when such analyses will be conducted on other surface epithelia. Whether keratinocytes undergo a ‘gradual model’ of differentiation from KRT14^+^ to KRT10^+^, or a ‘stepwise model’ wherein KRT14^+^/KRT10^+^ represents a transition state in keratinocyte differentiation, is an open issue that remains to be solved. Regardless, a higher resolution understanding of keratinocyte differentiation will provide significant insights in our understanding of skin homeostasis and disease.

The analyses reported here help shed new light on several open issues of high significance in the fields of keratin- and keratinocyte-related biology. First, how is the transcription of specific keratin type I and II genes, located amidts large keratin gene clusters located on two different chromosomes (chr. 12 and 17 in the human), so exquisitely coordinated as a function of the keratinocyte journey within the epidermis? The answer to this puzzle likely resides, at least in part, in the three dimensional organizational of chromosome domains, including the type I and type II keratin gene clusters, and the epidermal differentiation complex, in mammalian genomes (Botchkarev 2017). Second, how is a 1:1 molar ratio between type I and II keratin proteins achieved (Kim et al. 1984; Giudice and Fuchs 1987) under circumstances where individual keratinocytes show an imbalance between type I keratin transcripts, which are often dominant, and type II keratin transcripts? There is evidence that excess keratin of one type is inherently unstable and rapidly degraded (Giudice and Fuchs 1987; Kulesh et al. 1989) and also that keratin proteins are subject to ubiquination and proteasome-mediated degradation (Magin 1998; Ku 2000). Such post-transcriptional and post-tranlational layers of regulation are likely important since some keratins are seeminly not subject to a transcription-level match with a partner of the opposite type (e.g., *KRT2, KRT15* in epidermis). Related to this, the nature of the mechanisms that set individual keratin proteins to an optimal level in keratinocytes are also unknown. Again, here, pioneering studies in the field showed that type I and II keratins readily form hetero-oligomers in various types of epithelial cells (Hatzfeld and Franke 1985), adding to the complexity of this issue. Finally, one wonders about the significance and role, if any, of the mixing of progenitor-type (e.g., K5, K14, K15) and differentiation-type (e.g., K1, K10) keratins in hybrid keratinocytes at the time of their commitment to differentiation. A recent study provided strong evidence that K14-dependent disulfide bonding regulates a YAP1/Hippo-driven mechanism that gates entry of keratinocytes into differentiation within epidermis, with the key biochemical determinants conserved in some type I keratins (e.g, K10, K9) but not others (e.g., K15, K19) (Guo et al., 2020). Recent advances in spatial mRNA-seq (Marx 2021) further provide opportunities to observe transcriptional changes of keratinocytes as they initiate and undergo differentiation. Considering the spatially hierarchical organization of cell states within the epidermis, this method is poised to inform, and possibly revolutionize our understanding of emerging questions regarding keratinocytes differentiation in mammalian interfollicular epidermis. Thus, there are many properties and virtues awaiting to be discovered and further characterized in greater detail when it comes to keratin proteins, their regulation and roles in surface epithelia such as epidermis.

## Acknowledgments

The authors are very grateful to Dr. Raymond Cho (UCSF, USA) and Dr. Joseph Powell (Univ. Queensland, Australia) for making their single cell RNAseq data sets publicly available, and to members of the Coulombe laboratory for support. These efforts were supported in part by grant R01 AR079418 (to PAC) and grant T32 CA009676 (to CJR) from the National Institutes of Health. EC received support from the Center for Plasticity and Organ Design (CPOD) at the University of Michigan, and from the National Psoriasis Foundation.

