## Supplemental materials (Figs & Tables) for "Revisiting the significance of keratin expression in complex epithelia"

**List of contents**

**. Supplemental Methods**

**. Legends for Supplemental Figures and Tables**

Supplementary Fig. 1: Single cell RNA-seq analysis of healthy human trunk skin.

Supplementary Fig. 2: Additional analyses of scRNAseq data from human trunk skin.

Supplementary Fig. 3: Single cell RNA-seq analysis of healthy mouse ear skin.

Supplementary Fig. 4: Additional analyses of scRNAseq data from mouse ear skin.

Supplemental Table 1: Top 10 expressed genes in epidermal keratinocyte clusters.

Supplemental Table 2: Keratin expression levels across clusters.

Supplemental Table 3: Pairwise correlations across clusters.

Supplemental Table 4: Distribution of Custom G2M Scores across clusters.

#### Supplemental Methods

*Analysis of scRNAseq data from human trunk skin*

A standard “LogNormalize” (natural-log) Seurat pipeline (Hao et al. 2021) was applied for the re-analysis of a data set from Cheng et al. (2018). Cells with defined characteristics (between 500 and 7500 unique features; less than 15% mitochondrial content) were retained, yielding a total of 27,452 cells with informative content. Principal component analysis was applied for dimensional reduction, with the top 30 principal components retained for UMAP visualization (Suppl. Fig. 1A), construction of a shared nearest neighbor (SNN) graph, and subsequent clustering using the Louvain algorithm with a resolution setting of 0.5. This yielded twenty-one statistically distinct cell clusters (Suppl. Fig. 1A). Review of “top marker genes” and “most expressed genes” showed that fifteen clusters are comprised of epidermal keratinocytes, two correspond to appendageal keratinocytes, two correspond to melanocytes, one consists of Langerhans cells, and the last one is T cell-related (data not shown). Analyzing the entire array of cells partitioned in this fashion for specific marker genes such as *KRT14* (Suppl. Fig. 1B), *KRT10* (Suppl. Fig. 1C), *KRT2* (data not shown) and genes reflecting the S and G2M phased of the cell cycle (Suppl. Fig. 1, D and E) attests to a good balance between progenitor, actively dividing, and differentiating keratinocytes in this data set.

All analyzes of keratin expression were restricted to the fifteen clusters comprising epidermal keratinocytes, totaling 24,979 cells. Specifically, the six “Basal” keratinocyte clusters were defined by high abundance of *KRT5* and *KRT14*; the six spinous keratinocyte clusters defined by high abundance of *KRT1* and *KRT10*; and the three “Mitotic” clusters were characterized as keratinocytes based on high abundance of *KRT14*. The six clusters excluded from the keratinocyte analysis (and their markers) were “Langerhans” (high *CD74*), “T-Cell-like” (high *CXCR4*, *TRBC2*), two “Appendageal” (high *GJB2*, *TM4SF1*, *CYP1B1*, *KRT6A*), and two “Melanocytes” (high *DCT*, *PMEL*). Since our analyses were focused on the expression of keratins and not the clusters themselves, the latter were not reprocessed after the removal of the non-keratinocyte 6 clusters. It is worth noting, however, that data re-normalizing and re-clustering resulted in one fewer cluster where “Basal 1” and “Basal 6” were no longer independent, with all other cells largely assigned to the same orginal clusters (in this case, twelve principal components were used instead of fifteen to account for the decrease in information after removing four cell types).

*Analysis of scRNAseq data from mouse ear skin*

The same approach was used to reanalyze a data set from Lukowski et al. (2018), yielding nine statistically distinct cell clusters (Suppl. Fig. 3A). Adjustments made for lower cell counts and therefore lower dimensional data in this set included using the top 12 principal components and a resolution setting of 0.4 when clustering. Review of the “top marker genes” and “most expressed genes” showed that six clusters are comprised of epidermal keratinocytes (including a mitotic one), two are comprised of keratinocytes from epithelial appendages, and the last one is comprised of melanocytes (data not shown). Expression levels for *Krt14*, *Krt10*, and cell cycle genes are related in frames B, C, D and E of Suppl. Fig. 3. As with the human dataset the two “Basal” and one “Mitotic” clusters were defined by high expression of *KRT14* while the three “Spinous” clusters were defined by high *KRT1*. The “Melanocytes” and “Appendages” clusters respectively showed high *Dcn* and high *Defb6*, *Fst*, *Gstm5*, *Gsta3*. All analyzes of keratin expression in mouse skin were restricted to the six clusters comprising epidermal keratinocytes, totaling 4,761 cells.

*Composite scores and statistical analyses*

Composite scores were generated using a simple average of log expression values for a set of markers. For the “G2/M score” the gene markers used were *MKI67*, *TOP2A*, *CDC20*, *BUB3*, and *CCNB2*. For the “S Phase” score, the gene markers used were *RPA2, UHRF1, ATAD2, RFC2, and RRM2*. For the “Basal score” the gene markers used were *DST*, *POSTN*, and *COL17A1*. The Wilcoxon Rank Sum test was used for all top marker detection and differential expression analyses with tested genes limited to those detected at a minimum fraction of 0.25 in at least 1 of the clusters being compared to ignore genes less frequently expressed.

#### Legends

**Supplemental Figure 1. Single cell RNA-seq analysis of healthy human trunk skin.** (A-D) A data set published by Cheng et al. (2018) was reanalyzed using a standard Seurat pipeline (see Supplemental Material for details). UMAPs depicting (A) assigned Seurat clusters; (B) *KRT14* expression levels; (C) *KRT10* expression levels; (D) custom G2M Score and (E) custom S Phase Score across all cells in this data set (n= 27,452 cells with informative content, partitioned into 21 clusters).

**Supplemental Figure 2. Additional analyses of scRNAseq data from human trunk skin.** (A) Violin plot analysis of the ratio of *KRT14* / *KRT5* reads in Basal1-6 clusters (left), and the ratio of *KRT10* / *KRT1* reads in Spinous1-6 clusters (right). A dashed line across the Y-axis depicts a 1:1 balance between the type I and II keratin partners. (B, C) Correlations between specific type I and type II keratin genes across the entire array of epidermal keratinocytes (n= 24,979 cells). (B) *KRT2* vs *KRT10*. The red line reports on the linear correlation for all keratinocytes (n=24,979 cells). The magenta line reports on the linear correlation for Spinous6 keratinocytes (n= 408 cells). $R^2$ values are reported for each. (C) *KRT2* vs *KRT15,* which provides the equivalent of a negative control in this analysis of pairwais regulation in epidermis. (D) Heatmap relating expression of *POSTN*, *DST*, and *COL171A1* across the entire array of 24,979 keratinocytes. These three genes were selected to caculate a “Basal Score”, shown at right. See Suppl. Fig. 1 for source data and main text for additional details.

**Supplemental Figure 3. Single cell RNA-seq analysis of healthy mouse ear skin.**

(A-D) A data set published by Lukowski et al. (2018) was reanalyzed using a standard Seurat pipeline (see Supplemental Material for details). UMAPs depicting (A) assigned Seurat clusters; (B) *Krt14* expression levels; (C) *Krt10* expression levels; (D) custom G2M Score and (E) custom S Phase Score across all cells in this data set (n=5,446 cells with informative content, partitioned into 9 clusters).

**Supplemental Figure 4. Additional analyses of scRNAseq data from mouse ear skin.** (A) Heatmap relating expression of *Postn*, *Dst*, and *Col171a1* across the entire array of 4,761 mouse skin epidermal keratinocytes. These three genes were selected to caculate a “Basal Score”, shown at right. (B) Histogram reporting on the composite “Basal Score” across the population of mouse skin epidermal keratinocytes. See Suppl. Fig. 3 for source data and main text for additional details.

**Supplemental Table 1. Top 10 expressed genes in epidermal keratinocyte clusters.** Human trunk skin data (complement to Fig. 1B). See Suppl. Fig. 1 for details. Keratin genes are shown in red lettering.

**Supplemental Table 2. Keratin expression levels across clusters.**
Human trunk skin data (complement to Fig. 1B). See Suppl. Fig. 1 for details.

**Supplemental Table 3. Pairwise correlations across clusters.**
Human trunk skin data (complement to Fig. 1, C, D and E). See Suppl. Fig. 1 for details.

**Supplemental Table 4. Distribution of Custom G2M Scores across clusters.**
Human trunk skin data (complement to Fig. 2B) and mouse ear data (complement to Fig. 3E). See Suppl. Figs. 1 and 3 for details.

### Suppl. Figure 1

*Cohen, Johnson et al.*

*Human trunk skin data set*

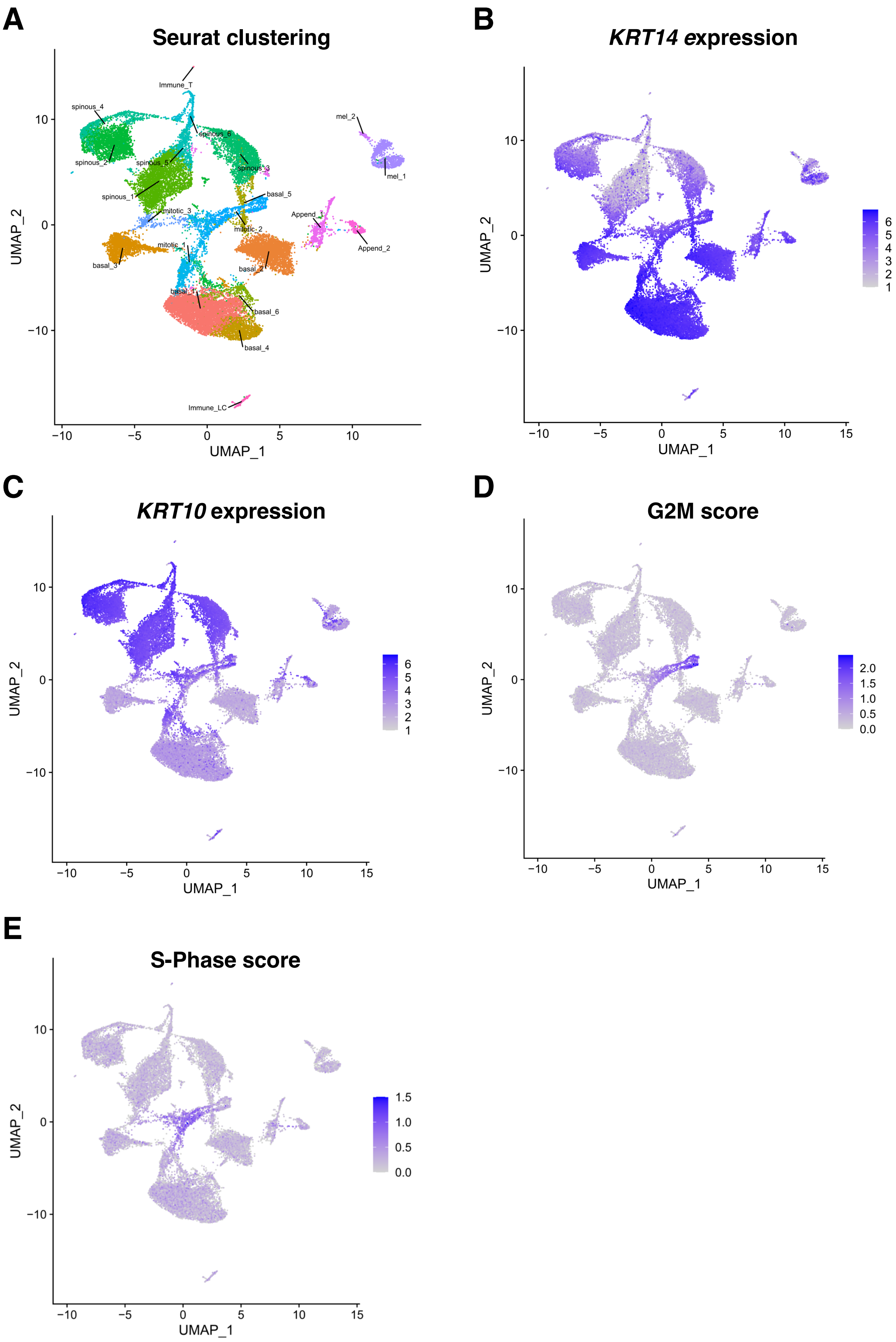

#### Suppl. Figure 2

*Cohen, Johnson et al.*

*Human trunk skin data set*

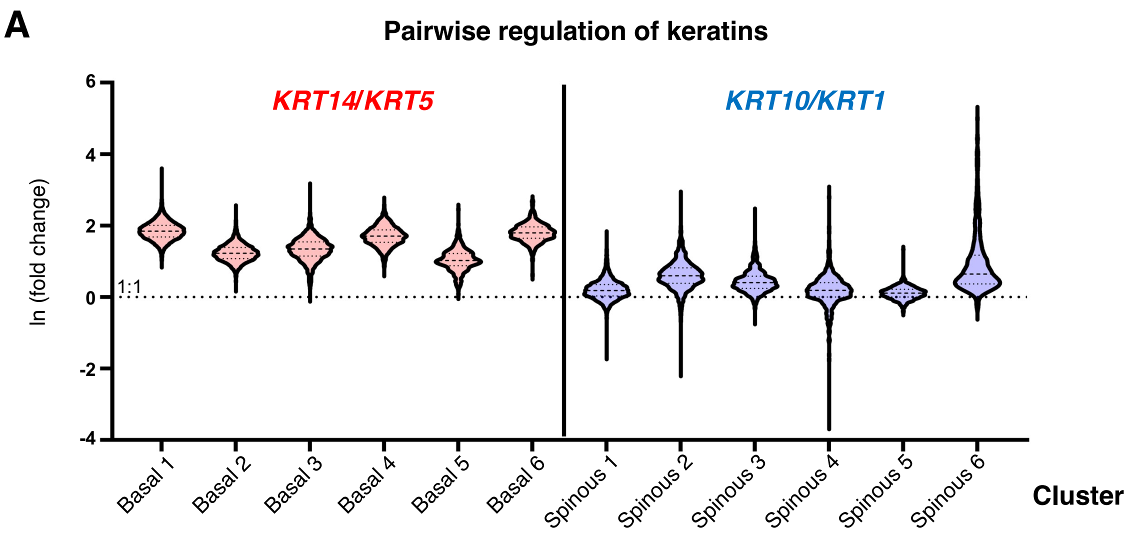


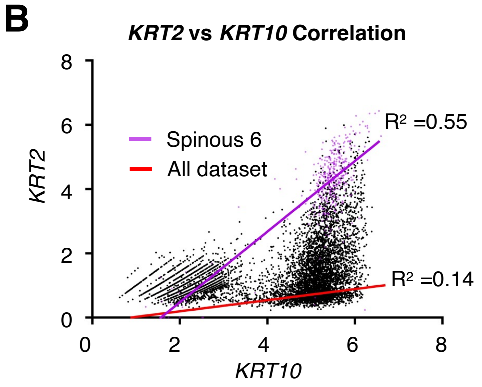


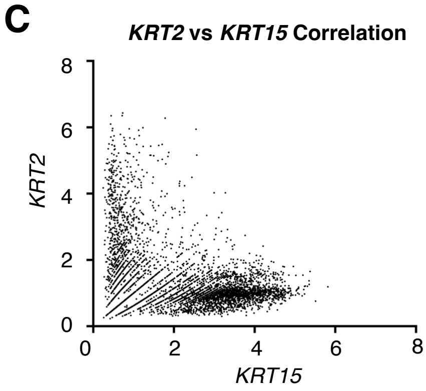


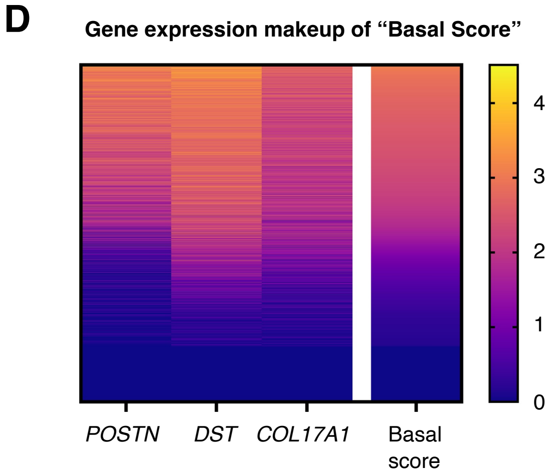

### Suppl. Figure 3

*Cohen, Johnson et al.*

*Mouse ear skin data set*

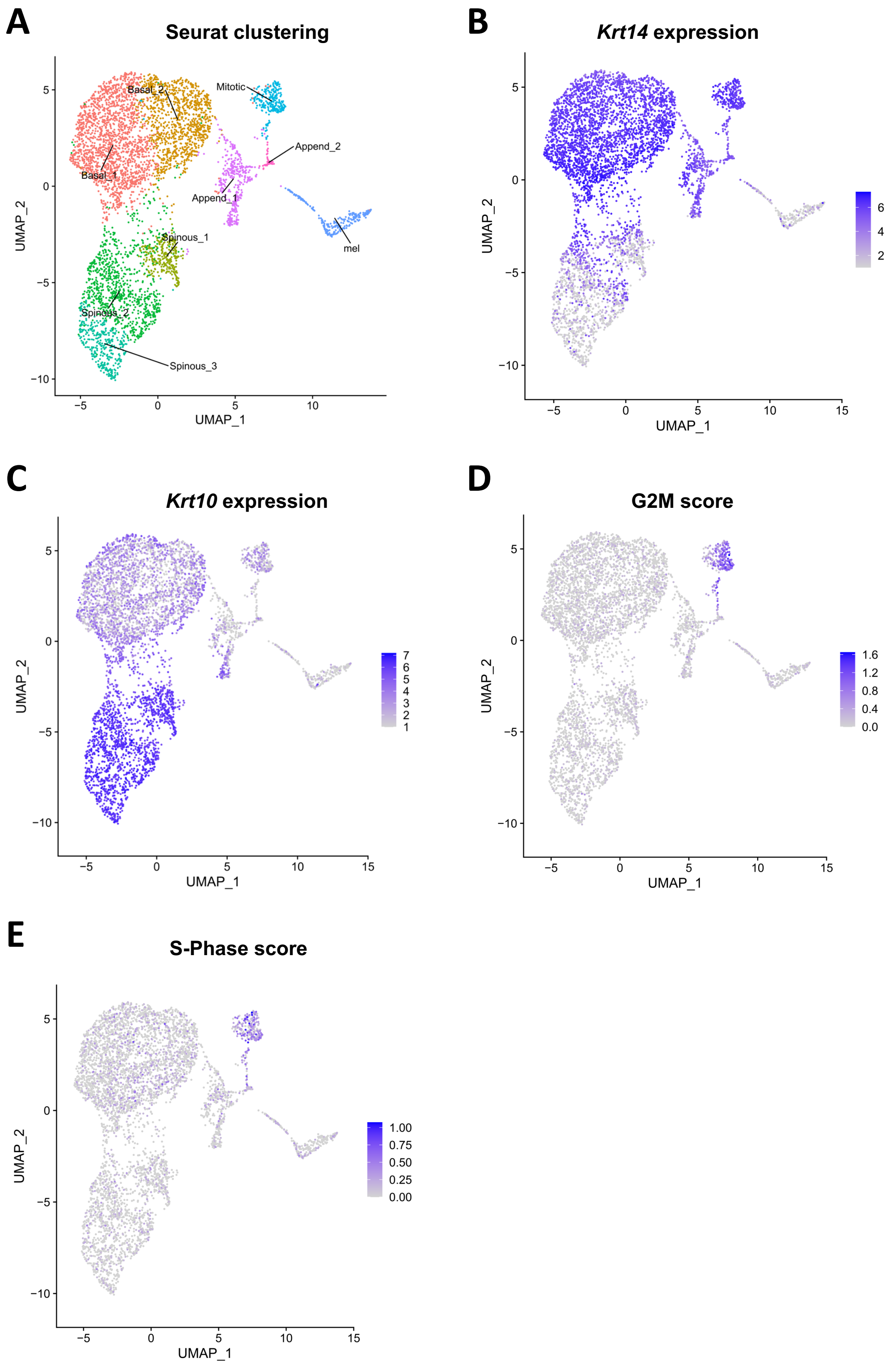

#### Suppl. Figure 4

*Cohen, Johnson et al.*

*Mouse ear skin data set*

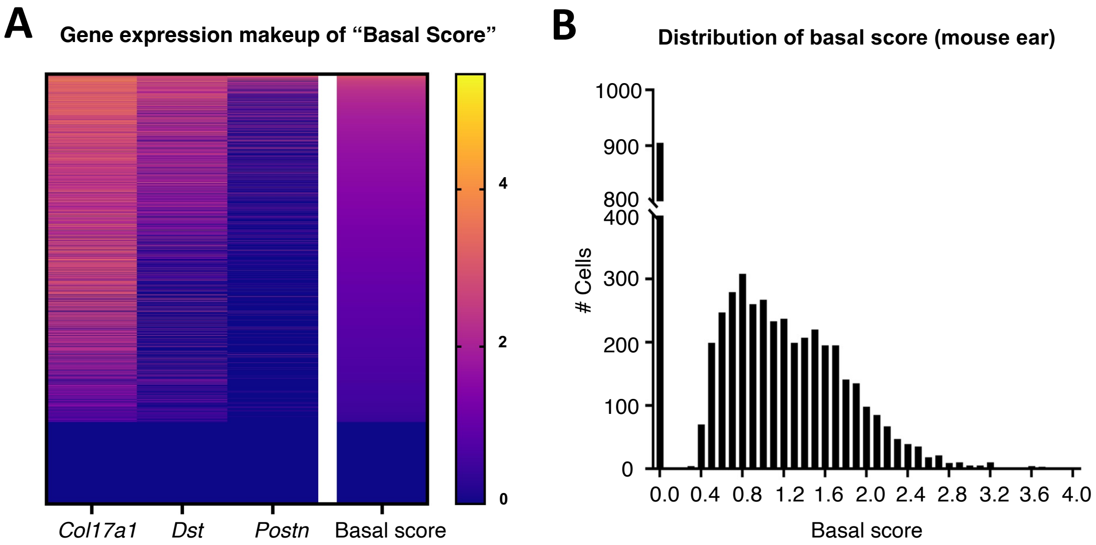

**Suppl. Table 1**

*Cohen, Johnson et al.*

#### Top 10 expressed genes per cluster (human trunk skin data)

##### Basal Clusters

| Rank | Basal 1 | Basal 2 | Basal 3 | Basal 4 | Basal 5 | Basal 6 |
|---|---|---|---|---|---|---|
| **1** | *MALAT1* | *MALAT1* | *MALAT1* | *MALAT1* | ***KRT14*** | *MALAT1* |
| **2** | ***KRT14*** | ***KRT14*** | ***KRT14*** | ***KRT14*** | *MALAT1* | ***KRT14*** |
| **3** | *EEF1A1* | *EEF1A1* | *EEF1A1* | *EEF1A1* | *EEF1A1* | *EEF1A1* |
| **4** | *CXCL14* | ***KRT5*** | ***KRT5*** | *CXCL14* | ***KRT5*** | *CXCL14* |
| **5** | ***KRT5*** | *CXCL14* | *CXCL14* | ***KRT15*** | *PTMA* | ***KRT5*** |
| **6** | *PTMA* | *PTMA* | *PTMA* | ***KRT5*** | *B2M* | *PTMA* |
| **7** | *TMSB4X* | *TPT1* | *B2M* | *PTMA* | ***KRT10*** | *TMSB4X* |
| **8** | *B2M* | *B2M* | *TPT1* | *TMSB4X* | *TPT1* | *B2M* |
| **9** | *PERP* | *TMSB4X* | *TMSB4X* | *B2M* | *MT1X* | *NFKBIA* |
| **10** | *H3F3B* | *H3F3B* | *JUN* | *H3F3B* | *DMKN* | *PERP* |

##### Mitotic Clusters

| Rank | Mitotic 1 | Mitotic 2 | Mitotic 3 |
|---|---|---|---|
| **1** | ***KRT14*** | *MALAT1* | ***KRT14*** |
| **2** | *MALAT1* | ***KRT14*** | *MALAT1* |
| **3** | *EEF1A1* | *PTMA* | ***KRT5*** |
| **4** | ***KRT5*** | *EEF1A1* | *EEF1A1* |
| **5** | ***KRT10*** | ***KRT5*** | *PTMA* |
| **6** | *PTMA* | *B2M* | *MT1X* |
| **7** | *B2M* | ***KRT10*** | *B2M* |
| **8** | *PERP* | *H3F3B* | *TPT1* |
| **9** | ***KRT1*** | *TMSB4X* | *PERP* |
| **10** | *DMKN* | *TUBA1B* | *DMKN* |

##### Spinous Clusters

| Rank | Spinous 1 | Spinous 2 | Spinous 3 | Spinous 4 | Spinous 5 | Spinous 6 |
|---|---|---|---|---|---|---|
| **1** | *MALAT1* | *MALAT1* | *MALAT1* | *MALAT1* | *MALAT1* | *MALAT1* |
| **2** | ***KRT10*** | ***KRT10*** | ***KRT10*** | ***KRT10*** | ***KRT10*** | ***KRT10*** |
| **3** | ***KRT1*** | *EEF1A1* | *EEF1A1* | ***KRT1*** | ***KRT1*** | *DMKN* |
| **4** | *EEF1A1* | ***KRT1*** | ***KRT1*** | *EEF1A1* | *DMKN* | ***KRT1*** |
| **5** | *DMKN* | ***KRT14*** | *DMKN* | *DMKN* | *EEF1A1* | *CALML5* |
| **6** | *B2M* | *DMKN* | *PTMA* | *PERP* | *KRTDAP* | ***KRT2*** |
| **7** | *PERP* | *B2M* | *B2M* | ***KRT14*** | *PERP* | *KRTDAP* |
| **8** | *PTMA* | *PERP* | *TPT1* | *B2M* | *PTMA* | *LOR* |
| **9** | *TPT1* | *PTMA* | *PERP* | *PTMA* | *B2M* | *EEF1A1* |
| **10** | *DSP* | *H3F3B* | *H3F3B* | *H3F3B* | *TMSB4X* | *PERP* |

#### Suppl. Table 2

*Cohen, Johnson et al.*

#### Keratin expression across clusters (human trunk skin data)

| | Basal Clusters | | | | | | Mitotic Clusters | | |
|---|---|---|---|---|---|---|---|---|---|
| Keratin | Basal 1 | Basal 2 | Basal 3 | Basal 4 | Basal 5 | Basal 6 | Mitotic 1 | Mitotic 2 | Mitotic 3 |
| ***KRT14*** | 6.2 | 5.8 | 5.8 | 6 | 5.7 | 6.1 | 5.8 | 5.2 | 5.7 |
| ***KRT5*** | 4.3 | 4.6 | 4.4 | 4.3 | 4.6 | 4.3 | 4.4 | 4.2 | 4.6 |
| ***KRT15*** | 2.8 | 2.7 | 3 | 4.3 | 1 | 3.1 | 0.9 | 1 | 0.5 |
| ***KRT10*** | 2.3 | 1.7 | 2 | 2.3 | 3.4 | 2.3 | 3.7 | 3.4 | 3.2 |
| ***KRT1*** | 1.7 | 1.2 | 1.5 | 1.7 | 3.2 | 1.6 | 3.3 | 2.7 | 3.2 |
| ***KRT2*** | 0.3 | 0.1 | 0.2 | 0.3 | 0.2 | 0.3 | 0.3 | 0.3 | 0.2 |

| | Spinous Clusters | | | | | |
|---|---|---|---|---|---|---|
| Keratin | Spinous 1 | Spinous 2 | Spinous 3 | Spinous 4 | Spinous 5 | Spinous 6 |
| ***KRT14*** | 2 | 3.9 | 3 | 3.3 | 1.4 | 1.8 |
| ***KRT5*** | 1.8 | 3.2 | 2.5 | 2.3 | 0.6 | 0.9 |
| ***KRT15*** | 0.2 | 0.7 | 0.3 | 0.7 | 0.2 | 0.3 |
| ***KRT10*** | 5 | 5.2 | 5 | 5.4 | 5.2 | 5.3 |
| ***KRT1*** | 4.8 | 4.6 | 4.5 | 5.2 | 5.1 | 4.4 |
| ***KRT2*** | 0.3 | 0.7 | 0.5 | 2.1 | 1.8 | 4.2 |

#### Suppl. Table 3

*Cohen, Johnson et al.*

#### Pairwise correlation across clusters (human trunk skin data)

##### Basal Clusters / Mitotic Clusters

| Pairwise $R^2$ | Basal 1 | Basal 2 | Basal 3 | Basal 4 | Basal 5 | Basal 6 | Mitotic 1 | Mitotic 2 | Mitotic 3 |
|---|---|---|---|---|---|---|---|---|---|
| ***KRT14-KRT5*** | 0.35 | 0.47 | 0.4 | 0.29 | 0.67 | 0.36 | 0.67 | 0.37 | 0.65 |
| ***KRT10-KRT1*** | 0.04 | 0.02 | 0.03 | 0.04 | 0.52 | 0.11 | 0.59 | 0.7 | 0.51 |
| ***KRT15-KRT5*** | 0 | 0.01 | 0 | 0.01 | 0.05 | 0.07 | 0.02 | 0.02 | 0.03 |
| ***KRT15-KRT14*** | 0.03 | 0.09 | 0 | 0.03 | 0.06 | 0.2 | 0.04 | 0.12 | 0.03 |
| ***KRT2-KRT10*** | 0 | 0 | 0 | 0 | 0.03 | 0.01 | 0.02 | 0.04 | 0 |

##### Spinous Clusters / Entire Dataset

| Pairwise $R^2$ | Spinous 1 | Spinous 2 | Spinous 3 | Spinous 4 | Spinous 5 | Spinous 6 | All cells |
|---|---|---|---|---|---|---|---|
| ***KRT14-KRT5*** | 0.45 | 0.44 | 0.5 | 0.54 | 0.32 | 0.68 | 0.8 |
| ***KRT10-KRT1*** | 0.46 | 0.42 | 0.51 | 0.82 | 0.65 | 0.35 | 0.84 |
| ***KRT15-KRT5*** | 0.05 | 0.04 | 0 | 0.15 | 0.13 | 0.3 | 0.33 |
| ***KRT15-KRT14*** | 0.15 | 0.17 | 0.01 | 0.31 | 0.2 | 0.35 | 0.48 |
| ***KRT2-KRT10*** | 0.01 | 0.01 | 0.07 | 0.08 | 0.03 | 0.56 | 0.14 |

#### Suppl. Table 4

*Cohen, Johnson et al.*

#### G2M score cluster distribution (human trunk skin data)

| Cluster | # Cells | % of G2M scoring cells |
|---|---|---|
| basal_1 | 1 | 0% |
| basal_3 | 1 | 0% |
| basal_4 | 1 | 0% |
| mitotic_1 | 8 | 1% |
| mitotic_2 | 714 | 97% |
| mitotic_3 | 1 | 0% |
| spinous_1 | 1 | 0% |
| spinous_2 | 1 | 0% |
| spinous_3 | 1 | 0% |
| spinous_4 | 3 | 0% |
| spinous_5 | 2 | 0% |
| spinous_6 | 1 | 0% |
| **Total** | **735** | |

#### G2M score cluster distribution (mouse ear skin data)

| Cluster | #Cells | % of G2M scoring Cells |
|---|---|---|
| basal_1 | 19 | 7% |
| basal_2 | 20 | 8% |
| mitotic | 215 | 81% |
| spinous_1 | 4 | 2% |
| spinous_2 | 8 | 3% |
| spinous_3 | 0 | 0% |
| **Total** | **266** | |
